# A Dataset for Deep Learning based Cleavage-stage Blastocyst Prediction with Time-lapse Images

**DOI:** 10.1101/2023.12.26.573382

**Authors:** Sijia Wang, Jing Fan, Hanhui Li, Mingpeng Zhao, Xuemei Li, David Yiu Leung Chan

## Abstract

Recent advances in deep learning and artificial intelligence techniques have obtained notable progress in automated embryo image analysis. However, most current research focuses on blastocyst-stage embryo evaluation (more than 5 days after in vitro fertilization), which may reduce the number of transferable embryos and increase the risk of canceled circles. Therefore, this paper aims to investigate the possibility of evaluating blastocyst development at the cleavage stage with deep neural networks (DNNs). To this end, we collect a dataset that consists of time-lapse images of more than 500 embryos (about 194k frames in total). We evaluate several widely used DNNs on the dataset, including those of single-frame architectures and multi-frame architectures. Experimental results show that the accuracy of different DNNs varies from 66.42% to 77.74% and we also provide the possible reasons behind the performance gap. Our dataset and code will be published soon to facilitate related research.

## 1 Introduction

The advent of time-lapse imaging technology in embryology has provided a wealth of information that can be utilized to improve the outcomes of in vitro fertilization (IVF) procedures. By continuously monitoring the development of embryos, time-lapse imaging allows for the detailed observation of morphokinetic parameters that are indicative of embryo viability [1]. The cleavage-stage blastocyst prediction is a critical aspect of this process, as it enables embryologists to select the embryos with the highest potential for successful implantation and subsequent development into a healthy fetus [2].

Recent advancements in artificial intelligence, particularly deep learning, have shown promise in enhancing the predictive capabilities of time-lapse imaging by automating the analysis of complex morphokinetic data (Tran et al., 2019). Deep learning algorithms, such as convolutional neural networks (CNNs), have been successfully applied to various medical imaging tasks, demonstrating their ability to learn from large datasets and identify patterns that may be imperceptible to the human eye [3].

However, the application of deep learning to cleavage-stage blastocyst prediction is contingent upon the availability of comprehensive and well-annotated datasets. Such datasets should include a diverse range of time-lapse images capturing the various stages of embryo development, along with corresponding outcomes data that indicate whether the embryos reached the blastocyst stage and resulted in a successful pregnancy [4]. The creation of a robust dataset is a challenging endeavor, as it requires the collection of images from multiple IVF centers to ensure variability and generalizability, as well as meticulous annotation by experienced embryologists.

In this research, we aim to construct and present a novel dataset specifically designed for the development and validation of deep learning models for cleavage-stage blastocyst prediction. The dataset comprises time-lapse images of preimplantation embryos, annotated with key developmental milestones and outcomes. We detail the methodology employed in the dataset’s creation, including the image acquisition process, the annotation protocol, and the steps taken to ensure data quality and integrity.

The significance of this dataset lies in its potential to facilitate the development of predictive models that can assist embryologists in making informed decisions regarding embryo selection. By leveraging deep learning, it is possible to automate the analysis of time-lapse images, thereby reducing the subjectivity and variability inherent in manual assessments [5]. Moreover, the predictive models derived from this dataset could lead to the identification of novel morphokinetic markers of embryo viability, further advancing the field of reproductive medicine [6].

The dataset is designed to be openly accessible to the research community, fostering collaboration and innovation in the application of deep learning to embryology. By providing a standardized resource for model development and benchmarking, we aim to accelerate the progress towards more accurate and reliable blastocyst prediction algorithms, ultimately improving IVF success rates and patient outcomes.

In conclusion, the creation of a dedicated dataset for deep learning-based cleavage-stage blastocyst prediction with time-lapse images represents a significant step forward in the intersection of reproductive medicine and artificial intelligence. This research not only contributes a valuable resource to the scientific community but also sets the stage for the development of advanced predictive tools that could revolutionize embryo selection practices in IVF clinics worldwide.

## 2 Related Work

The application of time-lapse imaging in assisted reproductive technology has been a subject of extensive research over the past decade. Time-lapse imaging allows for the continuous monitoring of embryos, providing a dynamic and detailed record of embryonic development, which has been shown to correlate with implantation potential [1]. The use of morphokinetic algorithms has been proposed to predict blastocyst formation and select embryos with the highest implantation potential [2].

In the realm of computational analysis, deep learning has emerged as a powerful tool for interpreting complex datasets. In medical imaging, deep learning, particularly convolutional neural networks (CNNs), has been successfully applied to tasks such as disease diagnosis, organ segmentation, and image classification (Litjens et al., 2017). In embryology, researchers have begun to explore the potential of deep learning to analyze time-lapse images and predict embryo viability [7].

Khan [5] discussed the potential of ‘big data’ in reproductive medicine, emphasizing the role of time-lapse imaging in generating large datasets that could be used to train deep learning models. The authors highlighted the need for robust datasets that are representative of the population and include various developmental stages and outcomes.

Jorgen [8] explored the use of deep learning for predicting IVF outcomes, demonstrating that 3D CNNs could be trained to predict embryo implantation from static images. It used multicentre data and known implantation results. However, their work also pointed out the private datasets and the need for publication, more diverse collections of annotated images to improve model performance.

Kragh [4] took a significant step by developing a deep learning approach for the real-time detection of early human embryos in time-lapse microscopy. Their work underscored the importance of high-quality datasets for training and validating deep learning models, as well as the challenges associated with annotating such datasets.

The creation of a specialized dataset for deep learning-based cleavage-stage blastocyst prediction with time-lapse images is a natural progression of these research efforts. Such a dataset would address the limitations identified in previous studies by providing a large, diverse, and well-annotated collection of time-lapse images that could be used to train more accurate and generalizable predictive models.

In summary, the related work in this field establishes a clear need for a large-scale, diverse, and well-annotated dataset for deep learning-based cleavage-stage blastocyst prediction. Such a dataset would not only advance the state of the art in embryo selection but also contribute to the broader field of reproductive medicine by providing a resource for developing more accurate and non-invasive predictive tools.

## 3 Data Collection

Embryo development images Collection

Collection period: August 1, 2021, to July 31, 2023

Collection sites: The Chinese University of Hong Kong Reproductive Center and Shenzhen Maternity & Child Healthcare Hospital Reproductive Center

Inclusion criteria:

1. Women undergoing IVF treatment.
2. Patients planning to use time-lapse imaging incubators for embryo culture.

Exclusion criteria: 1) Over half of the fertilized oocytes unable to divide correctly will be excluded. Fertilized oocytes unable to divide correctly are defined as having unclear imaging, significant blockage in the fertilized oocyte area, or over half of the fertilized oocyte area blocked or degraded. 2) Fertilized oocytes that are transplanted or cryopreserved before the 5th day will be considered inaccurately measured fertilized oocytes.

In this study, the incubator used for time-lapse imaging (TLI) is the Embryo Scope®. The time interval for TLI is set at 10 minutes. The CO2 concentration is set to 6.0%, and the temperature is maintained at 37.0°C. For conventional embryo culture, G-TL medium (Swiss Vitrolife company) is used, while G-IVF medium (Swiss Vitrolife company) is used for the culture of immature oocytes.

## 4 Methodology

Extensive architectures have been proposed since the era of deep learning and it is impractical to evaluate all of them. Considering morphological and morphokinetic features are the mainstream criteria in embryo evaluation, we mainly consider two paradigms for blastocyst prediction with time-lapse images, as shown in Figure 3: Given a sequence of embryo images, we can utilize (i) single-frame networks that take the last frame of the image sequence as input, or (ii) multi-frame networks that use the whole sequence. Evaluating single-frame networks allows us to investigate the effects of different network architectures in modeling morphological features, while multi-frame networks can reveal the importance of temporal information (morphokinetic features).

**Figure 1:**
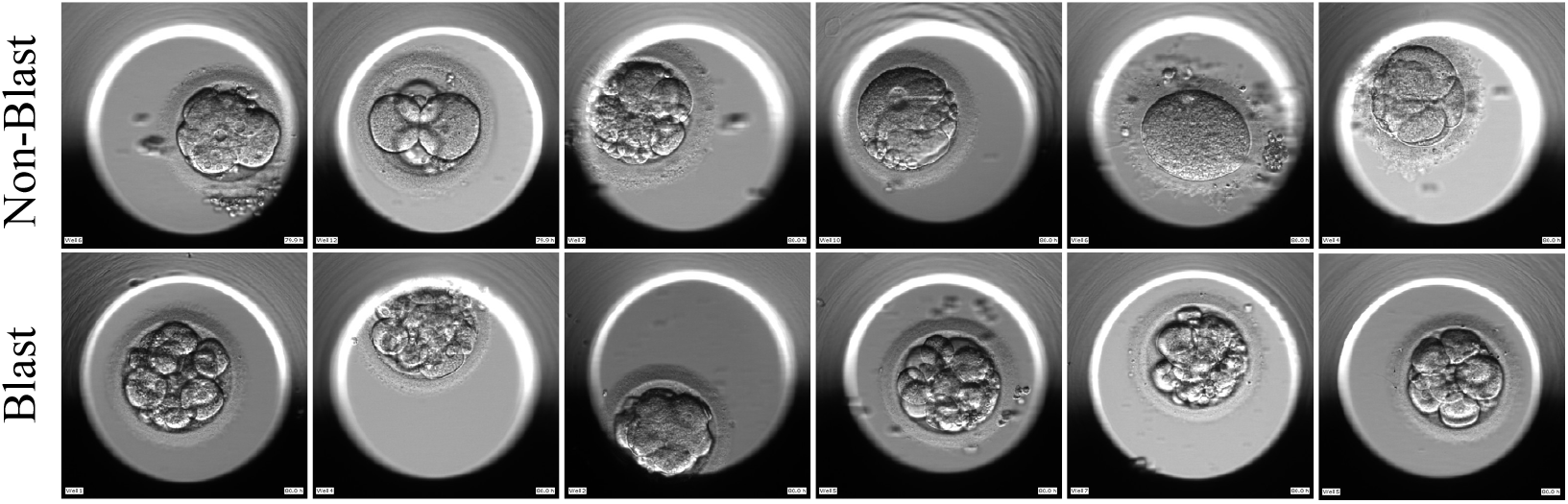
Visual examples of embryos arrested before blastocyst formation (top row) and developed into blastocysts (bottom row).

**Figure 2:**
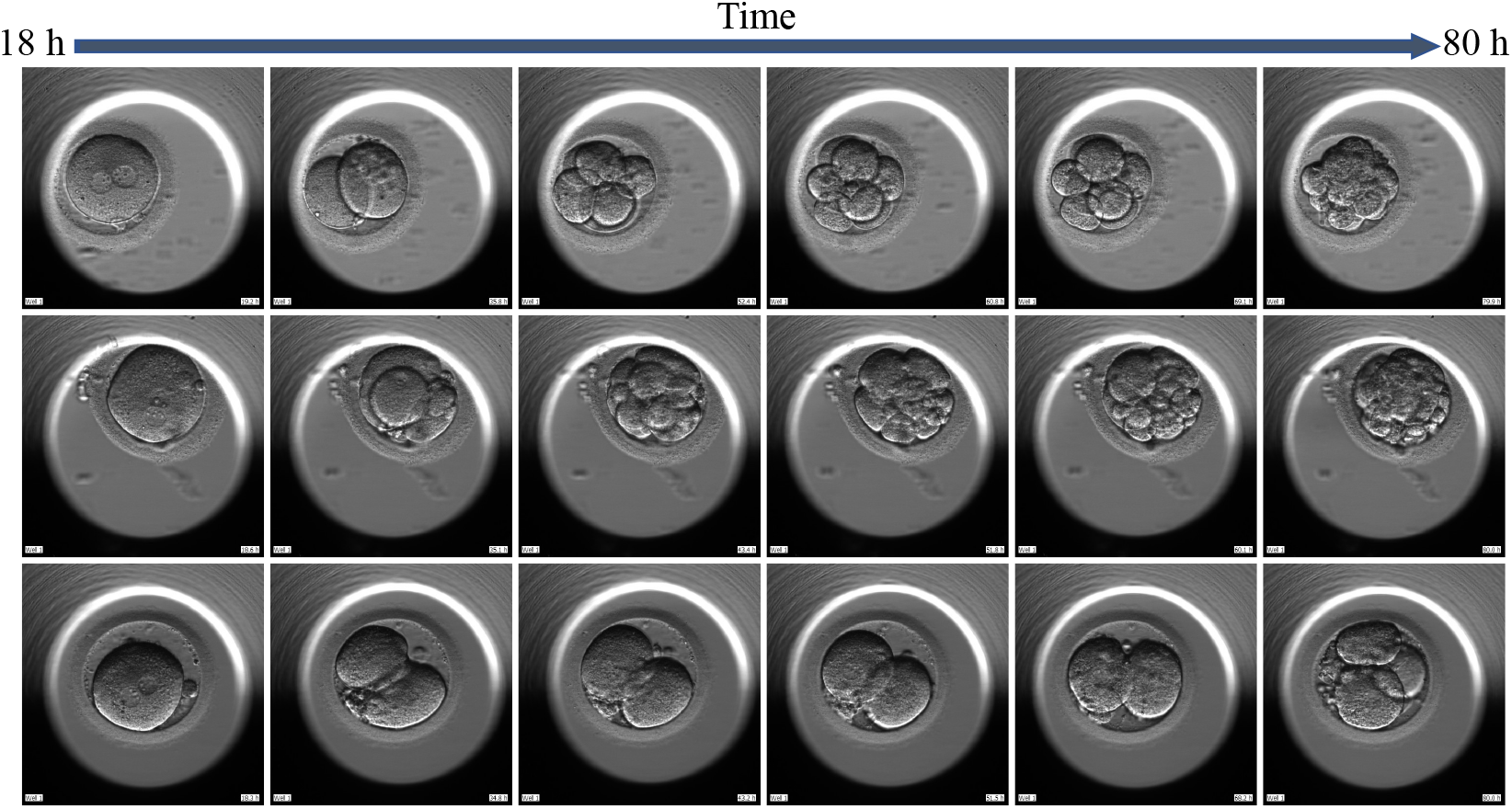
Examples of time-lapse sequences collected in the proposed dataset.

**Figure 3:**
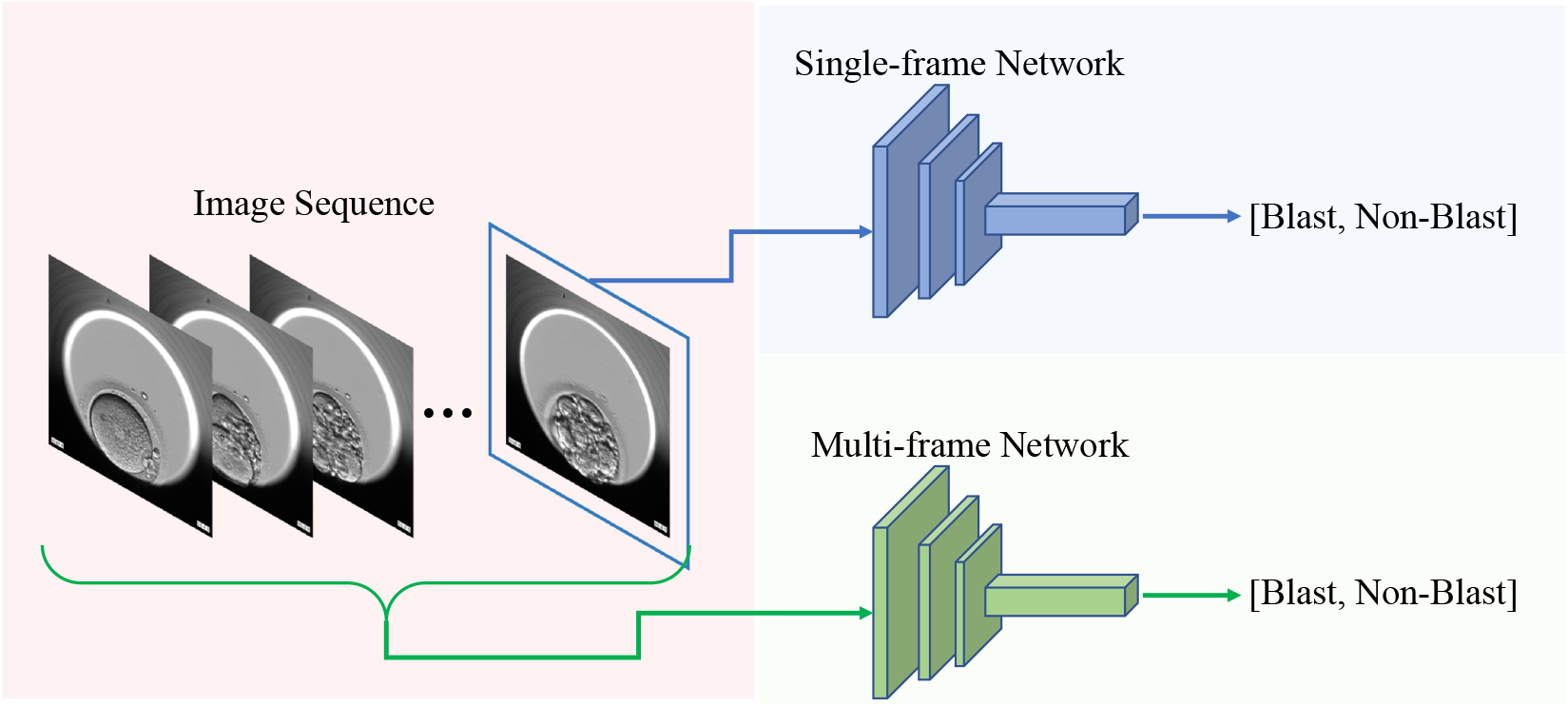
Paradigms for early embryo prediction considered in this paper.

In both paradigms, we formulate blastocyst development prediction as a binary classification task that predicts whether an embryo is arrested before blastocyst formation (denoted as non-blast) or develops into a blastocyst (denoted as blast). We consider minimizing the cross-entropy loss between the predicted binary label and the ground truth as the learning objective of all networks.

### Setup

Our experiments are conducted within the PyTorch framework, using two GeForce RTX 2080 Ti graphics cards. To construct a class-balanced training/test split, we construct the training set by randomly selecting half embryos from the sets of blasts and non-blasts, and use the remaining embryos for testing. Multiple data augmentation methods are used to prevent over-fitting, including color jitter, random cropping, rotation, and horizontal flipping. All networks are trained with the AdamW optimizer [9] with a fixed learning rate of 10^*−*3^ and a weight decay factor of 10^*−*5^. As the computational cost of multi-frame networks is usually higher than that of single-frame networks, we set different batch sizes, training epochs, and image sizes for these two paradigms. As summarized in

Table 2, we train single-frame networks with all images resized to 400 × 400, a batch size of 32 for 500 epochs, and train multi-frame networks with an image size of 224 × 224 a batch size of 16 for 100 epochs.

**Table 1:** Statistics of the proposed benchmark.

|  |  |
| --- | --- |
| Number of Patients | 83 |
| Number of Frames | 194,713 |
| Number of Embryos | 531 |
| Blast : Non-Blast | 264 : 267 |
| Frame interval | 5 mins |
| Phases | 0 h ~ 80 h |

**Table 2:** Setup of experiments.

|  | Single-frame | Multi-frame |
| --- | --- | --- |
| Batch Size | 32 | 16 |
| Optimizer | AdamW [9] |  |
| Learning Rate | $1e^{-3}$ | |
| Weight Decay | $1e^{-5}$ | |
| Epochs | 500 | 100 |
| Input Frame | 1 | 32 |
| Frame Size | $400 \times 400$ | $224 \times 224$ |

#### 4.1 Single-frame Networks

We mainly evaluate canonical single-frame networks that are provided in the Torchvision library, including ResNet series [10, 11, 12], MobileNet [13], Inception [14], DenseNet [15], and vision transformers [16]. Here we briefly introduce these networks and explain why they are chosen.

##### ResNet

ResNet is one of the most widely used convolutional neural networks. It proposes the strategies of residual connections and identity mapping to achieve the training of deep network architectures. We evaluate ResNets with different numbers of layers, e.g., ResNet-18 denotes the ResNet with 18 layers.

##### Wide-ResNet

Instead of depth, Wide-ResNet emphasizes the importance of the width of networks, i.e., the number of channels. As time-lapse images exhibit fewer patterns compared with natural images, we select Wide-ResNet as one of our baselines to investigate whether a wide network still has significant advantages in this case.

##### Inception

Inception networks combine convolution filters of different sizes into a convolutional block to better model multi-scale features. This baseline is used to validate if small morphological features help to predict blastocyst development. We adopt InceptionV3 in our experiments.

##### ResNeXt

ResNeXt combines the advantages of ResNet and Inception by adopting multiple branches in the residual learning block.

##### DenseNet

DenseNet proposes a dense connection strategy that connects all the precedent features of a dense block. This strategy can exploit features of various levels effectively. We include this baseline to investigate the effects of image features from different layers.

##### MobileNet

MobileNet is designed for edge computing scenarios where network inference cost is a major consideration for applications. This baseline is considered as we are interested in finding whether complicated architectures or high-capacity networks are necessary for blastocyst prediction.

##### ViT

Except for the above CNN-based architectures, we also consider cutting-edge vision transformers (ViT) due to their notable performance in related tasks. The core of ViTs is self-attention modules that capture long-range feature correlations, yet they also result in expensive computational costs.

#### 4.2 Multi-frame Networks

The core of multi-frame networks is to aggregate features of multiple frames and explore temporal information. Hence in this section, we use the backbone of ResNet-50 as our per-frame image encoder unless specified, and combine it with different temporal models and feature aggregation strategies. Specifically, we consider three groups of temporal models, including (i) LSTM [17] and GRU [18], which are canonical sequential models that merge features frame by frame; (ii) Mixtures of 2D convolutions and 3D convolutions (including I3D [19], ResNet3D, MC and R(2+1)D [20]). (iii) Multi-frame networks (MFNet [21]) that adopt more tailored architectures instead of the ResNet backbone. The details of these models are listed as follows:

##### LSTM

An LSTM unit typically consists of blocks for long-term and short-term memory that are used to tackle the vanishing gradient problem in sequential modeling. Given the features of multiple frames, these memory blocks utilize multiple gates to select important features.

##### GRU

GRU merges the gates in LSTM units and hence provides a more compact and efficient architecture. To adopt LSTM and GRU, we first extract per-frame feature maps with the ResNet backbone. We then conduct bidirectional feature map propagation with the corresponding sequential model, i.e., from the first frame to the last one and reversely. We concatenate the last feature maps in the forward and backward propagation process, so that they can be utilized for prediction as in single-frame networks.

##### ResNet3D

We construct ResNet3D by replacing all 2D convolutions in ResNet with 3D convolutions.

##### MC

MC denotes the simple mixture of 2D and 3D convolutions. Particularly, we construct an MC network that adds multiple 3D convolutional blocks after the 2D backbone to process concatenated multi-frame feature maps.

##### R(2+1)D

The idea of the R(2+1)D architecture is to separate a 3D convolution into a spatial 2D convolutional block followed by a temporal 1D convolutional block.

##### I3D

Similar to ResNet3D, I3D inflates the Inception architecture via changing 2D blocks like convolutions and poolings to their 3D counterparts.

##### MFNet

MFNet aims at improving the efficiency of multi-frame networks. It introduces a multi-fiber architecture that divides convolution filters into groups to reduce the number of parameters and also encourages mutual feature refinements among groups. Similar to MobileNet, we select MFNet to investigate whether complicated architectures are necessary for our task.

## 5 Results

### Performance of single-frame networks

Table 3 reports the experimental results of single-frame networks. We can see that the accuracy of sing-frame networks varies from 66.42% to 74.72% and is not proportional to network complexity. For example, ViT-large has 302.78M parameters, yet its accuracy is similar to that of ResNet50, which has 23.56M parameters only. Similarly, Wide-ResNet101 has 124.84M parameters but its performance is lower than that of the ResNet101 baseline. The major reason behind this phenomenon is that large-capacity networks require more training data, which is hard to satisfy in our case.

**Table 3:** Performance of single-image based classification networks.

| Method | Parameters (M) | Accuracy |
| --- | --- | --- |
| MobileNetV3 [13] | 4.20 | 66.79 |
| InceptionV3 [14] | 21.79 | 72.08 |
| ResNet18 [10] | 11.68 | 70.94 |
| ResNet50 [10] | 23.56 | 69.43 |
| ResNet101 [10] | 42.50 | 67.92 |
| ResNet152 [10] | 58.14 | 69.06 |
| ResNeXt50 [11] | 22.98 | 70.19 |
| ResNeXt101 [11] | 86.74 | 66.42 |
| Wide-ResNet50 [12] | 66.83 | 67.17 |
| Wide-ResNet101 [12] | 124.84 | 66.42 |
| Dense-121 [15] | 6.95 | 73.58 |
| Dense-169 [15] | 12.48 | 74.72 |
| ViT-base [16] | 85.40 | 70.57 |
| ViT-large [16] | 302.78 | 69.81 |

Moreover, we notice that the accuracies of DenseNet series are significantly higher than other baselines. Note that the core of DenseNets is to maintain and combine features of multiple levels, especially for those that are low-level. This indicates that certain low-level features are more likely to carry important morphological information and hence should be leveraged to tackle blastocyst prediction.

### Performance of multi-frame networks

The accuracies of our multi-frame baselines are summarized in Table 4. From these results, we can observe that the performance of multi-frame networks is consistently better than that of single-frame networks. This is reasonable, as multi-frame networks can better leverage morphokinematic information in TLI sequences. ResNet-R(2+1)D achieves the highest accuracy (77.74%) and its complexity is moderate, compared with a few large single-frame baselines. This suggests that with limited network capacity, exploring morphokinematic information might be more beneficial for our task, compared with that is morphological.

**Table 4:** Performance of multi-frame networks.

| Method | Parameters (M) | Accuracy |
| --- | --- | --- |
| ResNet-LSTM [17] | 22.19 | 73.96 |
| ResNet-GRU [18] | 19.56 | 75.47 |
| I3D [19] | 25.84 | 72.08 |
| ResNet3D [20] | 33.15 | 73.96 |
| ResNet-MC [20] | 11.47 | 75.09 |
| ResNet-R(2+1)D [20] | 31.30 | 77.74 |
| MFNet [21] | 7.69 | 73.96 |

## 6 Conclusion and Future Work

We delve into the implications of our experimental findings and draw overarching conclusions regarding the application of deep learning models in predicting cleavage-stage embryo development based on time-lapse images.

The experimental setup involved the use of the PyTorch framework. To ensure a balanced training/testing set, half of the dataset, comprising randomly selected embryos from both blastocyst formation and non-blastocyst formation categories, was used for training, while the remaining half was used for testing. Various data augmentation techniques, such as color jittering, random cropping, rotation, and horizontal flipping, were employed to mitigate overfitting.

Two distinct network architectures, namely single-frame and multi-frame networks, were utilized in the experiments. Single-frame networks included classic architectures from the Torchvision library, such as ResNet, MobileNet, Inception, DenseNet, and Vision Transformers. On the other hand, multi-frame networks primarily featured ResNet-50 as the main frame image encoder, coupled with diverse temporal models and feature aggregation strategies, including LSTM, GRU, I3D, ResNet3D, MC, R(2+1)D, and MFNet.

The detailed performance analysis of single-frame and multi-frame networks revealed intriguing insights. Single-frame networks exhibited accuracies ranging from 66.42% to 74.72%, showcasing a disproportionate relationship between network complexity and performance. In contrast, multi-frame networks achieved accuracies below 77.74%, with relatively minor variations in performance across different models. Notably, the ViT model, with a substantial parameter count of 302.78 million, demonstrated accuracy similar to the more lightweight ResNet50, which had only 23.56 million parameters.

These findings prompt several noteworthy discussions. Firstly, the observed performance of single-frame networks suggests that, in the context of cleavage-stage embryo prediction, increasing model complexity may not necessarily lead to significant improvements in accuracy. This raises questions about the need for intricate architectures in scenarios where simpler models suffice.

Secondly, the performance of multi-frame networks, while achieving higher accuracies, presents a relatively small margin of improvement compared to single-frame counterparts. The similarities in accuracy between models with vastly different parameter counts, such as ViT-large and ResNet50, highlight the importance of exploring efficient model architectures that balance accuracy and computational cost.

The discussion extends to the implications for practical applications in assisted reproductive technology (ART). While our experiments provide valuable insights into the potential of deep learning for early-stage embryo assessment, further research is necessary to validate these findings on diverse datasets and clinical scenarios. The translation of these models into real-world clinical practice requires addressing challenges related to interpretability, generalizability, and ethical considerations.

In conclusion, our study contributes to the evolving field of automated embryo assessment by investigating the use of deep neural networks for predicting blastocyst embryo development. The findings underscore the importance of carefully balancing model complexity and computational efficiency, with potential implications for the broader application of deep learning in reproductive medicine. As we move forward, collaborative efforts between researchers, clinicians, and technologists will be crucial to harness the full potential of artificial intelligence in improving outcomes in assisted reproduction.

## 7 Acknowledgment

This work was supported by Shenzhen-Hong Kong-Macau S&T Program (Type C) (SGDX2020110309500101).

